# Quantifying cortical maturational aspects during different vigilance states in preterm infants by advanced EEG analysis

**DOI:** 10.1101/2025.05.14.653972

**Authors:** Gaia Burlando, Sara Uccella, Valentina Marazzotta, Sheng H Wang, J Matias Palva, Monica Roascio, Andrea Rossi, Luca Antonio Ramenghi, Lino Nobili, Gabriele Arnulfo

## Abstract

Preterm birth is associated with numerous neurodevelopmental adverse outcomes, even in the absence of acquired lesions, as it occurs during a critical period of brain development. Clear organization of vigilance states can be recognized from 30–32 weeks postmenstrual age (PMA). In this study, we investigated whether spatial and temporal properties of neuronal oscillatory dynamics (*i.e.*, phase synchronization, bistability, and cross-frequency coupling) during different vigilance states provide insights into cortical maturation in preterm infants born very low birth weight (VLBW) at low neurological risk and devoid of detectable brain lesions.

We analyzed artifact-free video-polysomnographic data from 11 VLBW preterm infants (PMA at recording: 33.0 ± 1.6 weeks) who underwent brain MRI at term-equivalent age. For each vigilance state, we computed the weighted Phase Lag Index (wPLI), Bistability Index (BiS), and Phase-Amplitude Coupling (PAC), both globally and across anterior and posterior regions, and examined their correlation with PMA at recording.

wPLI, BiS, and PAC showed specific trends across vigilance states. BiS and PAC exhibited posterior-to-anterior differences and correlated with PMA.

Our study suggests that these electrophysiological markers, particularly BiS and PAC, may serve as indices to monitor aspects of cortical maturation in VLBW at low neurological risk.

## Introduction

Preterm birth, defined as delivery before 37 weeks of gestation, occurs in approximately 10% of live births worldwide and is associated with a heightened risk of adverse neurodevelopmental outcomes, even in the absence of overt brain injury (Marlow et al. 2021; Perin et al. 2022; Uccella et al. 2023; Malova et al. 2024). This is because preterm birth interrupts a critical phase of brain development that would otherwise take place in utero, particularly during the third trimester, when processes such as neurogenesis, neuronal migration, synaptogenesis, and the establishment of cortico-cortical and cortico-subcortical connections are at their peak (McQuillen and Ferriero 2005; Uylings 2006; Dubois et al. 2014; Uccella et al. 2024). During this sensitive developmental window, cortical and subcortical structures undergo progressive maturation and integration: long-range and short-range networks, including intra- and inter-hemispheric pathways, strengthen along distinctive gradients (*e.g.*, posterior-to-anterior), and these changes are reflected in the brain’s emerging functional activity (Ramenghi et al. 2007; Thomason et al. 2013, 2014; Jakab et al. 2014; van den Heuvel and Thomason 2016). Accordingly, the early postnatal period in preterm newborns represents a crucial window to evaluate and refine methods for assessing how brain maturation advances outside the womb, especially when preterm birth occurs at the earliest gestational ages. Quantitative methods for assessing brain activity at this early stage are essential to identify early biomarkers of neurodevelopment and to understand their potential correlation with long-term outcomes.

Electrophysiological activity, as captured through EEG, reflects these maturational processes and can serve as a non-invasive readout of developing network function. However, electrophysiological brain activity is neither static nor homogeneous: it fluctuates according to vigilance states, particularly sleep, which is the dominant condition during this period of life. In fact, sleep occupies more than two-thirds of total time in preterm neonates (Roffwarg et al. 1966; Uccella et al. 2025) and is thought to actively contribute to brain maturation (Frank 2017; Blumberg et al. 2022). Importantly, sleep itself is not uniform, as it comprises different stages, such as active sleep (AS) and quiet sleep (QS), each associated with distinct patterns of neuronal oscillations and brain connectivity (Vanhatalo and Kaila 2006; Shiraki et al. 2024). These stages emerge progressively and follow specific developmental trajectories, paralleling the maturation of underlying neural circuits (Grigg-Damberger 2016; Blumberg et al. 2020; Uccella et al. 2024). For example, AS, considered a precursor to rapid eye movement (REM) sleep, is prominent in early life and in neonates, particularly in preterms, frequently appears at sleep onset, while QS, associated with non-REM features, gradually increases with age (Stern et al. 1969; DeHaan et al. 1977). The close relationship between electrophysiological patterns, the organization of behavioral states, and neurodevelopment suggests that studying EEG features across different vigilance states could yield critical insights into brain maturation. Crucially, each state reflects specific configurations of cortical-subcortical interactions and connectivity dynamics. To capture the underlying principles governing these dynamic changes, we adopted a systems-level approach grounded in the synchrony-criticality framework, which posits that the brain operates near a critical regime between hypo- and hyper-synchronization, a balanced condition that supports both stability and adaptability (Haldeman and Beggs 2005; Chialvo 2010; Cocchi et al. 2017). In this state, brain activity exhibits hallmarks of scale-free systems, including bistability and non-linear transitions, enabling efficient information processing (Kinouchi and Copelli 2006; Lotfi et al. 2021). Dysregulation of this balance has been implicated in developmental and pathological conditions, including epilepsy (Wang et al. 2023, 2024; Burlando et al. 2025). Based on this framework, we focused our analyses on three EEG-derived features that reflect key aspects of network dynamics: a) phase synchronization, coordination of oscillatory activity across regions (Buzsáki and Draguhn 2004; Fries 2005; Palva et al. 2005, 2018); b) bistability, the brain’s tendency to shift between more and less synchronized states (Freyer et al. 2009; Wang et al. 2023); c) cross-frequency coupling, hierarchical coordination of activity across timescales (Canolty and Knight 2010; Palva and Palva 2018; Siebenhühner et al. 2020). These features offer a multiscale view of developing cortical dynamics and are particularly sensitive to both maturational processes and behavioral states.

This study aims to examine how neuronal synchrony, bistability, and cross-frequency coupling vary with postmenstrual age (PMA) and across sleep-wake states in a cohort of neurologically normal preterm infants without brain lesions. By quantifying how EEG markers evolve with PMA and how they differ across vigilance states, we aimed to capture the underlying architecture and dynamics of early cortical maturation. We hypothesized that frontal regions would exhibit greater bistability and excitability due to delayed maturation compared to posterior areas and that such spatial gradients would correlate with PMA independently of behavioral state. The ultimate goal of this work is to establish normative, state-specific developmental trajectories of key EEG biomarkers in preterm infants at low neurological risk. These biomarkers could serve as early quantitative developmental markers that, in the future, may help identify infants at risk for later neurodevelopmental disorders. Given the heterogeneity of outcomes in preterm populations and the lack of specific risk factors for neurodevelopmental disorders other than intellectual disability, such biomarkers of evolution are crucial for guiding early interventions.

## Materials and Methods

### Patients and data acquisition

This observational study was part of a larger cohort study on preterm infants born very low birth weight (VLBW) approved by the Local Ethical Review Board (CERL number approval 0028224/21 of 6/10/2021). It was conducted in accordance with the Declaration of Helsinki. All data were recorded at the Neonatal Intensive Care Unit of the Gaslini Children’s Hospital. Parents provided informed consent for the study. Subjects were recorded once they were hemodynamically stable and free from sedation. The study included only VLBW infants who underwent a prolonged full-channel polysomnography between 30 and 36 weeks of PMA, allowing us to assess maturational changes during this critical period in a sufficient period of artifact-free recording. Additionally, participants had a normal neurological examination and a brain MRI free from any congenital or anatomically detectable brain lesions at term-equivalent age.

Eleven VLBW preterm infants (7 females) were included in the study, born between 28 and 35 weeks of gestation (31.0 ± 2.2 weeks, mean ± SD), with appropriate birthweight and head circumference for their gestational age (Bertino et al. 2010). The PMA at the recording session was 33.0 ± 1.6 weeks. The sample size aligns with previous EEG studies on preterm neonates’ vigilance states.

### Signal preprocessing

All subjects underwent a video-polysomnography recording using a Brain QUICK® (Micromed) system. Scalp cortical activity was measured using a full-band EEG amplifier (FbEEG, SD Plus Flexi Clinic Micromed). We used non-polarizable Sintered Ag/AgCl cup electrodes (5 mm diameter, 19 mm² recording area) and secured them with electroconductive paste. Once the right and left parasagittal and lateral longitudinal lines were outlined, eight active surface electrodes were placed according to the modified 10/20 system for neonates. The reference electrode was placed on Fz, away from the bregmatic fontanelle, and the ground electrode on Pz.

The data were then selected to retain 10 monopolar EEG channels, excluding EMG, ECG, and EOG, acquiring bandpass signals (0.15–134 Hz) from these channels based on the 10-20 International System, at a sampling rate of 512 Hz.

We excluded all harmonics of 50-Hz line noise using a series of notch FIR filters with a 1 Hz band-stop width. We employed a bipolar referencing scheme by pairing neighboring electrode contacts while excluding a common reference (Fig. 1a). After this step, the channels for each subject encompassed the following bipolar derivations: Fp1–C3, C3–O1, Fp1–T3, T3–O1, T3–C3, Fp2–C4, C4–O2, Fp2–T4, T4–O2, C4–T4, Fp1–Fp2, C3–C4, O1–O2. The EEG recordings were manually scored by clinicians with over five years of experience in neonatal sleep. Quiet wakefulness (QW), sleep onset active sleep (SOAS), active sleep (AS), and quiet sleep (QS) were identified and segmented into 30-second epochs, as recommended by the American Academy of Sleep Medicine (Grigg-Damberger 2016). The best minutes were carefully selected, and epochs were extracted only if they were free of artifacts (mean ± SD: 8.1 ± 1.6 min for QW; 7.9 ± 1.7 min for SOAS; 7.8 ± 2.0 min for AS; and 12.0 ± 2.3 min for QS).

**Figure 1:**
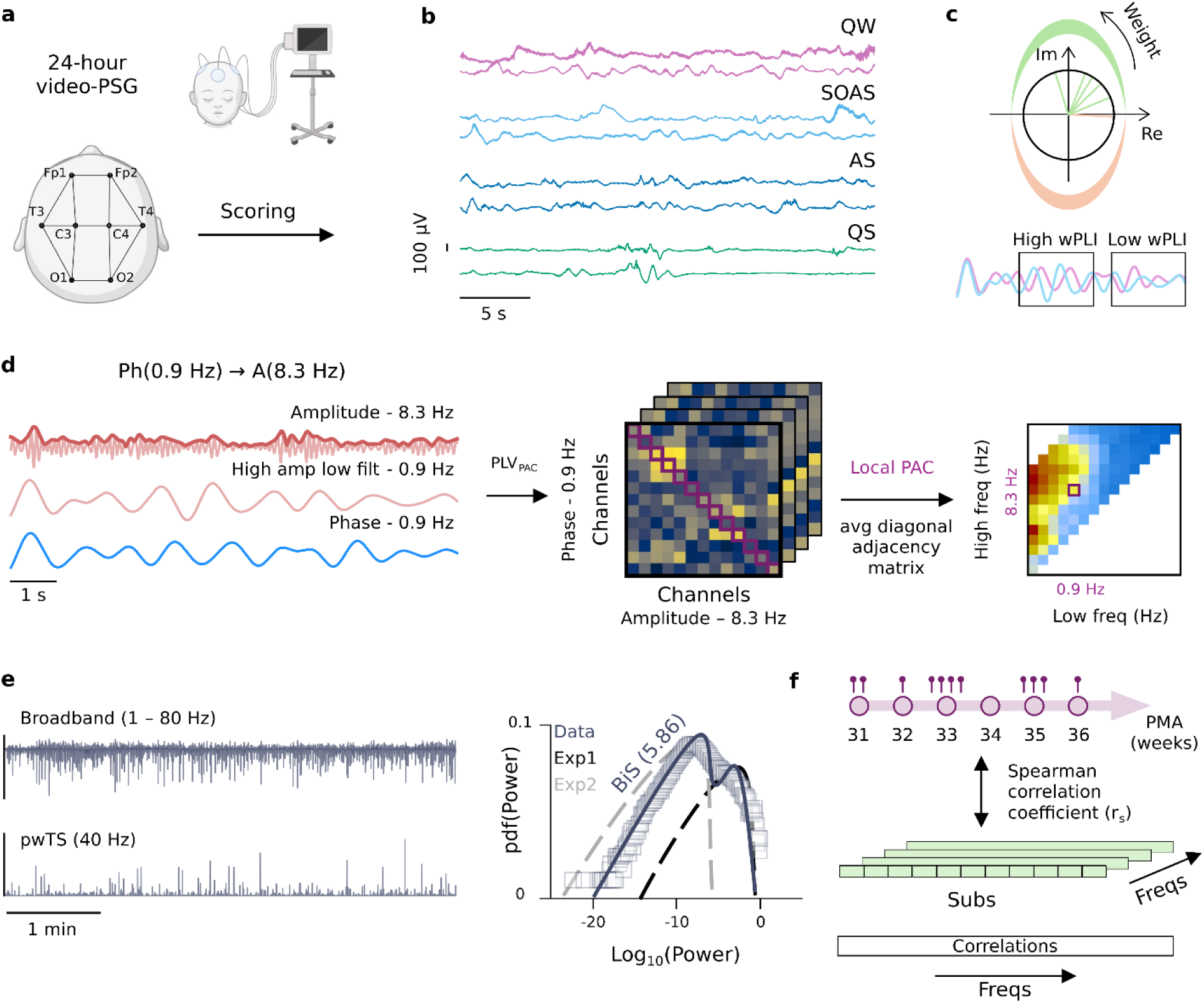
Study design and analysis overview. **(a)** EEG recordings were collected from a cohort of very preterm infants during sleep and quiet wakefulness. Following artifact rejection procedures, a bipolar montage was employed for subsequent analysis. **(b)** The recordings were classified into various behavioral states: quiet wakefulness (QW), sleep onset active sleep (SOAS), active sleep (AS), and quiet sleep (QS). **(c)** Illustration of the weighted phase-lag index (wPLI) (orange: phase lead; green: phase lag). Example of two time series initially showing high wPLI (phase synchronized), followed by a period of low wPLI. **(d)** Illustration of Phase-Amplitude Coupling (PAC) Analysis: A broadband signal is filtered using a Morlet wavelet with a low frequency (LF) of 0.9 Hz, while another signal is filtered with a high frequency (HF) of 8.3 Hz. The envelope of the HF signal is extracted and processed with a Morlet wavelet at the same LF as before. The Phase Locking Value (PLV) is then calculated between the two time series, resulting in subject-specific adjacency matrices for all behavioral states. Local PAC is computed for all the channels and averaged to obtain a single value for each low-frequency-high-frequency pair. **(e)** Left: 5 minutes of broadband and 40-Hz narrow-band power time series. Right: The fitting for the bistability index (BiS). **(f)** Spearman correlations between PMA and wPLI, BiS, or nPAC are computed for each subject across all frequencies, yielding a spectrum of correlation coefficients.

### Spectral power estimates

To evaluate the power spectral characteristics of EEG signals across the four behavioral states in preterm infants, we calculated the power spectral density (PSD) for all bipolar channels and states in each subject using Welch’s method, applied within the frequency range of 0.25 to 40 Hz with a resolution of 0.25 Hz. The power spectrum was computed using a Hamming window, and then the spectra were averaged across subjects within the same sleep stage.

### Phase synchrony estimates

We estimated interareal phase interactions at the individual subject level using the weighted phase-lag index (wPLI) (Vinck et al. 2011). This measure is computed between signals from different EEG channels and is often used to help mitigate the potential influence of volume conduction in neuronal oscillation analysis (Vinck et al. 2011; Palva et al. 2018). WPLI ranges from 0 to 1, where a higher value indicates stronger phase synchronization, implying that the phases of the two signals tend to align more consistently over time. First, we applied a time-frequency decomposition by Morlet-wavelet filtering the data into narrow-band time series for 50 frequencies, ranging from 0.2 Hz to 40 Hz (n° cycles = 7.5), logarithmically spaced (Tallon-Baudry et al. 1996). Then, wPLI was computed as:

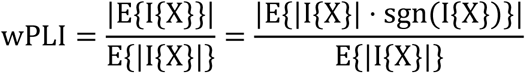

where I{X} represents the imaginary part and X is the cross-spectrum between the two EEG time series.

### Cross-frequency coupling of slow and fast rhythms

Cross-frequency interaction was assessed by calculating phase-amplitude coupling (PAC). We quantified PAC using the phase locking value (PLV), which measures the consistency of the phase relationship between two frequency bands across different time points, providing an estimate of the coupling strength (Vanhatalo et al. 2004). We measured the PLV to investigate the modulation of higher-frequency (HF) oscillation amplitudes by the phase of lower-frequency (LF) oscillations, separately for each behavioral state. The strength of PAC was quantified as:

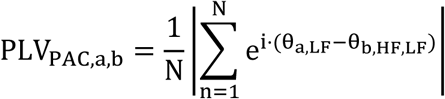

where θ_b,HF,LF_ is the phase of the amplitude envelope of the HF signal filtered with a Morlet filter at LF. Local PAC was obtained where a = b.

As low frequencies, we used the same 50 as in the phase synchrony estimates, while the high frequencies were computed using 19 slow-to-fast ratios ranging from 1:2 to 1:100. The PAC matrices, initially structured in a low frequency-to-ratio format, were interpolated and mapped into a low frequency-to-high frequency space to improve readability.

Results show normalized PAC (nPAC), where nPAC = PLV_PAC,observed_/PLV_PAC,surrogate_, so that nPAC > 1 indicates PAC above the null hypothesis level.

### Bistability of neuronal oscillations estimates

We estimated the level of bistability within a narrow-band neuronal oscillation by computing the bistability index (BiS) (Freyer et al. 2011; Wang et al. 2023). The process involves fitting the observed probability distribution of the narrow-band power time series with both a single exponential and a bi-exponential model. Subsequently, BiS is determined by comparing these two models using the Bayesian information criterion. A BiS close to zero means that the single-exponential model is a more likely model for the observed time series, whereas, for a BiS greater than 0, the most likely model for the observed time series is the bi-exponential model. We applied Morlet wavelet filtering to extract narrow-band time series for the same 50 frequencies as before and then computed BiS separately for each vigilance state.

### Statistical analysis

Statistical analyses were performed using Python 3.11. When comparing metrics across the four vigilance states, we used the Kruskal-Wallis test followed by Benjamini-Hochberg (BH) correction for multiple comparisons along the frequency axis. In this case, to assess the effect size for comparisons involving more than two groups (i.e., for the four vigilance states), we computed the η^2^ as follows:

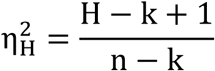

where H is the value obtained in the Kruskal-Wallis test, k is the number of groups (i.e., the four vigilance states), and n is the total number of observations. The eta-squared ranges from 0 to 1, with values typically classified as: 0.01 ≤ η^2^ ≤ 0.06 (small effect), 0.06 ≤ η^2^ ≤ 0.14 (moderate effect), and η^2^ ≥ 0.14 (large effect) (Cohen 2008).

### Spatial maturational distribution

For pairwise comparisons (*e.g.*, posterior vs. frontal derivations), we applied the Wilcoxon rank-sum test, also with BH correction. To quantify the effect size for these two-group comparisons, we calculated Cohen’s d using the formula:

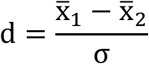

where x̄_1,2_ are the means of the two samples, and σ is the pooled standard deviation. Cohen’s d was interpreted as follows: 0.2 ≤ d ≤ 0.5 (small effect), 0.5 ≤ d ≤ 0.8 (medium effect), and d ≥ 0.8 (large effect) (Cohen 2013). Correlation analyses were performed using Spearman’s rank correlation coefficient. Statistical significance was defined as *p* < 0.05 after BH correction.

### Correlation estimates with PMA

To explore the dynamic changes in neuronal networks driven by cortical maturation, we analyzed the correlations between phase synchronization, bistability, and cross-frequency coupling with PMA. To assess the predictive value of BiS, wPLI, and nPAC on PMA, we employed three separate linear mixed-effects models (LMMs) (Raudenbush and Bryk 2002). These models were fitted using restricted maximum likelihood estimation (REML), with each metric as the dependent variable and PMA, sex, and sleep stage (SOAS as the reference stage) as fixed effects. Frequency bands were incorporated as random effects to account for repeated measures.

## Results

### Vigilance states distinctly modulate global neuronal oscillatory dynamics

To examine the impact of vigilance states on neuronal oscillatory dynamics, we analyzed power density, phase synchronization, bistability, and cross-frequency coupling across states.

Amplitude of local neuronal oscillations was affected by vigilance states, with QS showing the lowest power, followed by SOAS and AS, while QW exhibited the highest (*p* < 0.05, Kruskal-Wallis test, Benjamini-Hochberg corrected) (Fig. 2a). These differences spanned the entire frequency range (0.2–40 Hz), except in the θ–α band (6–10 Hz), and with stronger effect size (η^2^ > 0.14) in the slow-wave band (< 1Hz) and β-γ band (> 20Hz). At a large-scale level, neuronal oscillations seemed to interact in a state- and frequency-specific manner at this early stage of life. Spectral analysis of global phase synchronization revealed distinct wPLI peaks across different vigilance states (Fig. 2b). During QS, wPLI was significantly higher than in the other vigilance states in the δ band (2–3 Hz), while QW, SOAS, and AS showed higher wPLI in the θ band (4–6 Hz) (*p* < 0.05, Kruskal-Wallis test, BH uncorrected).

**Figure 2:**
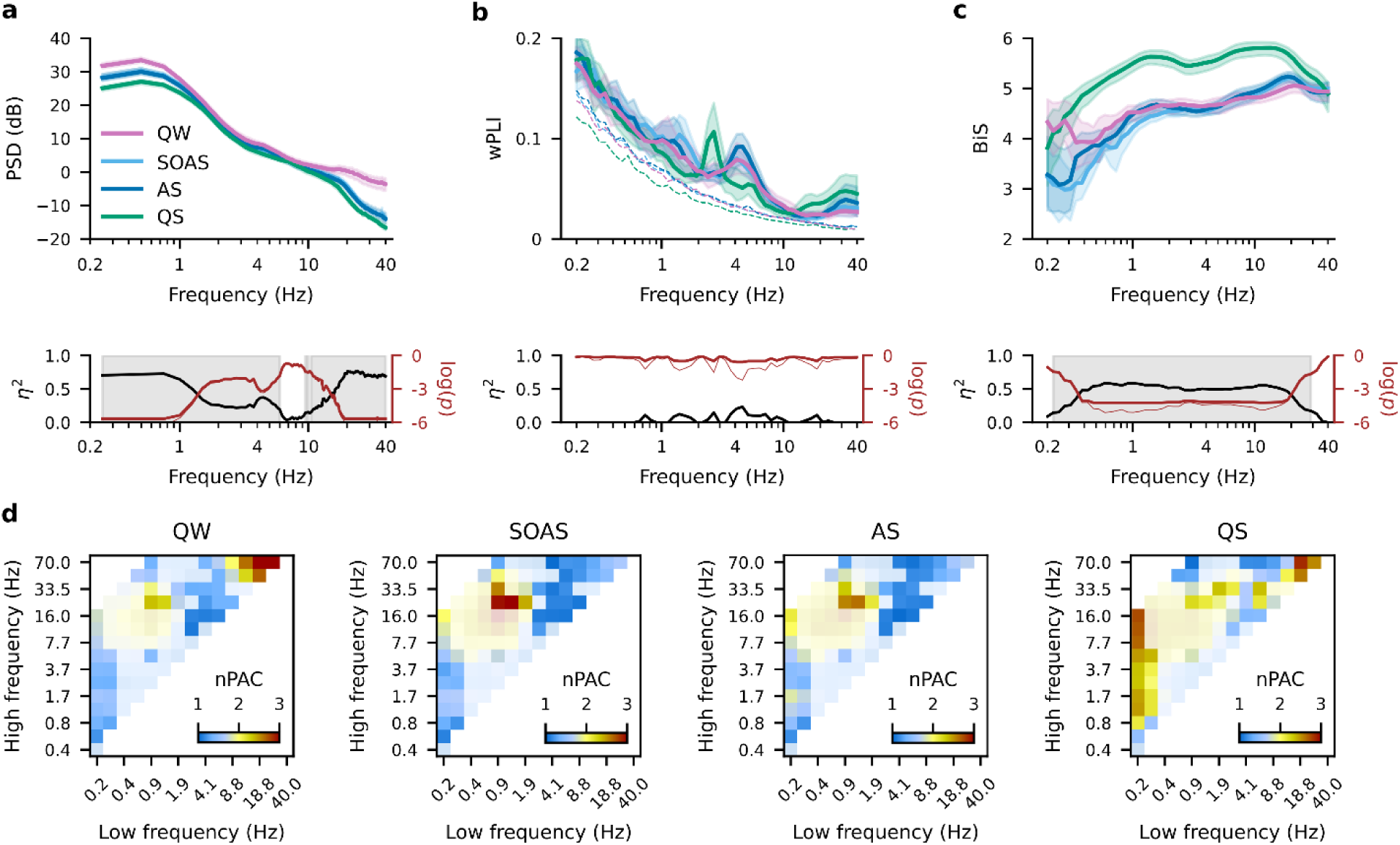
Neuronal dynamics are modulated by different vigilance states. **(a)** Power spectral density (PSD), **(b)** global weighted phase-lag index (wPLI), and **(c)** global bistability index (BiS) across the different behavioral states. The shaded areas represent the variability computed from 100 bootstraps (95^th^ percentile). The second row shows the spectra of effect size (η^2^, black) and *p*-value (reddish) for each frequency. The *p*-value is represented as a thin line (uncorrected) and a thick line (BH-corrected). Grey areas indicate η^2^ > 0.14 (large effect) and *p* < 0.05 after Benjamini-Hochberg FDR correction. **(d)** Cohort-averaged normalized phase-amplitude coupling (nPAC) for each behavioral state. Warmer colors indicate stronger coupling, while cooler colors represent weaker coupling. nPAC matrices are alpha-shaded after Kruskal-Wallis test and Benjamini-Hochberg FDR correction. Regions with less transparency correspond to areas with η^2^ > 0.14 (large effect) and BH-corrected *p*-value < 0.05.

Differences extended beyond amplitude modulation of single oscillations as we observed a significant difference in global bistability across vigilance states (*p* < 0.05, Kruskal-Wallis test, BH corrected) (Fig. 2c). BiS values during QS were higher compared to other vigilance states across the frequency range of 0.2–30 Hz, with the strongest effect (η^2^ > 0.14) in the 0.5–12 Hz. SOAS and AS showed similarly lower BiS values in the slow-oscillations band (< 0.5 Hz) compared to the other vigilance states. BiS values during QW in the very low frequency range were higher with respect to AS and SOAS.

Cross-frequency coupling also revealed peculiar frequency modulations (Fig. 2d). Quiet wakefulness displayed higher coupling between the phase of high-β oscillations (20–30 Hz) and γ amplitudes (55–70 Hz). In SOAS and AS, the phase of δ-band oscillations was coupled with β-oscillation amplitudes. In contrast, QS showed a more widespread area of modulation between phases < 1 Hz and broad-band oscillations (1–16 Hz) and between the phase of β oscillations (20–30 Hz) and γ amplitudes (55–70 Hz).

Altogether, these results revealed that: QW was characterized by the highest amplitudes in a wide range of frequencies, a broadband mild level of bistability, prominent peaks in θ and β-to-γ band wPLI, and increased coupling between β and γ oscillations; SOAS and AS had similar characteristics with mid-level amplitudes, a reduced bistability of slow oscillations, a peak in θ band phase synchronization, and significant effect of slow-wave oscillations in modulating β-γ band amplitudes; finally, QS had the smallest amplitude, a significantly higher bistability across a broad range of frequencies, a prominent peak in wPLI at around 2 Hz, and significantly higher modulation between slow-oscillation phases and the amplitudes of faster rhythms.

### Neuronal dynamics reflect spatial differences in posterior and frontal derivations

Next, we analyzed whether neuronal dynamics differ between mid-posterior (C3–O1, T3–O1, C4–O2, T4–O2, O1–O2) and mid-frontal (Fp1–C3, Fp1–T3, Fp2–C4, Fp2–T4, Fp1–Fp2) derivations. In large-scale connectivity analysis, we observed elevated wPLI levels in posterior derivations for frequencies < 1 Hz, during QW, SOAS, and AS (*p* < 0.05, Wilcoxon ranksums test, BH uncorrected) (Fig. 3a). Posterior derivations exhibited lower levels of wPLI during QW (*p* < 0.05, Wilcoxon ranksums test, BH corrected) and SOAS (*p* < 0.05, Wilcoxon ranksums test, BH uncorrected) in the 5–10 Hz range. Additionally, we observed a peak in wPLI in frontal derivations at 1 Hz during SOAS (*p* < 0.05, Wilcoxon ranksums test, BH uncorrected), absent in posterior derivations. No significant differences between posterior and frontal derivations were observed in QS.

**Figure 3:**
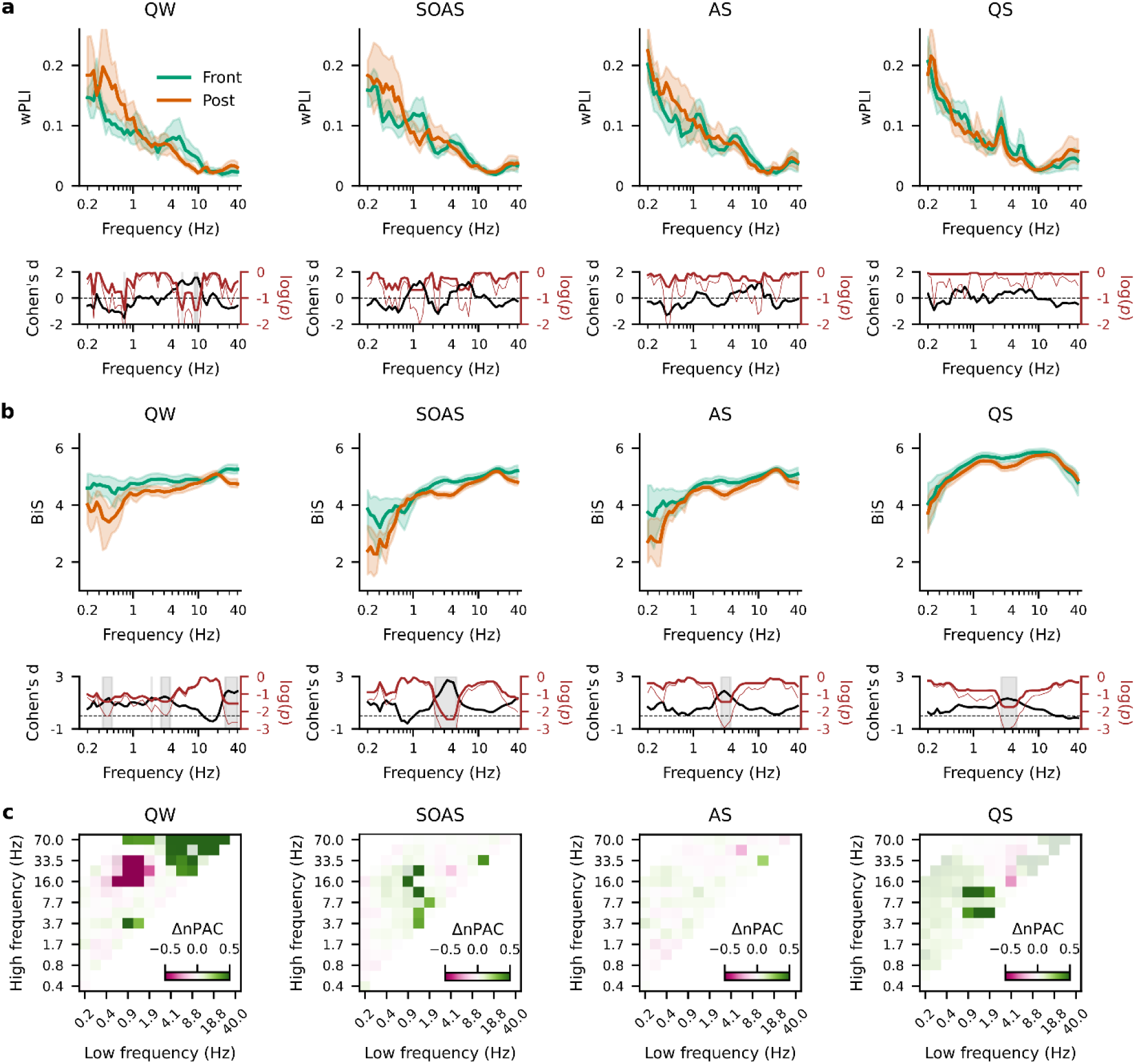
Neuronal dynamics differ from posterior to frontal derivations. **(a)** Comparison between frontal vs. posterior weighted phase-lag index (wPLI) and **(b)** bistability index (BiS), separately for each behavioral state. The shaded areas represent the variability computed from 100 bootstraps (95^th^ percentile). The second row shows the spectra of effect size (Cohen’s d, black) and *p*-value (reddish) for each frequency. The *p*-value is represented as a thin line (uncorrected) and a thick line (BH-corrected). Grey areas indicate d > 0.8 (large effect) and *p* < 0.05 after Benjamini-Hochberg FDR correction. **(c)** Difference between frontal and posterior normalized phase-amplitude coupling (nPAC) for each behavioral state. Green hues indicate nPAC_Frontal_ > nPAC_Posterior_, whereas pink hues indicate the opposite. Alpha shading was applied after pairwise Wilcoxon ranksums test and Benjamini-Hochberg FDR correction. Regions with less transparency correspond to areas with d > 0.8 (large effect) and uncorrected *p* < 0.05.

Posterior derivations showed significantly lower values of BiS in the δ band (2–4 Hz) than anterior derivations across all the vigilance states (*p* < 0.05, Wilcoxon ranksums test, BH corrected) (Fig. 3b). Moreover, quiet wakefulness was characterized by significantly lower BiS values in the posterior compared to anterior derivations in the low-frequency range (< 1 Hz) and in β–γ bands (25–40 Hz) (*p* < 0.05, Wilcoxon ranksums test, BH corrected). In QW, posterior derivations showed stronger δ-to-β/γ coupling, while frontal derivations exhibited stronger θ-to-β and γ coupling (p < 0.05, uncorrected). Similarly, in SOAS and QS, stronger frontal modulation emerged between δ and θ-to-α bands. However, none of these PAC differences remained significant after correction for multiple comparisons (Fig. 3c).

### Bistability and local phase-amplitude coupling decrease with age in preterm infants

We first examined the relationships of each metric (wPLI, BiS, and local nPAC) with PMA, vigilance state, and frequency as random factors separately (Tab. 1).

**Table 1:** Summary of the linear mixed-effects model assessing the relationship between each dependent variable (wPLI, BiS, and nPAC), postmenstrual age (PMA), vigilance state, and sex. The table reports the estimated coefficients (Coefficient), standard errors (Std. Err.), z-scores (z), p-values (P > |z|), and 95% confidence intervals ([0.025, 0.975], (L) and (U), respectively) for each predictor. The intercept represents the expected dependent variable value for the reference category (SOAS for vigilance states; female for sex). The coefficients for ‘Stage’ indicate the differences in the dependent variable relative to SOAS, while ‘Sex [T.M]’ represents the difference between males and females. ‘Group Var.’ denotes the variance of the random effect (frequency bands). Statistically significant results (*p* < 0.05) are marked with an asterisk (*).

| **a) Dependent Variable: wPLI** |  |  |  |  |  |  |
| --- | --- | --- | --- | --- | --- | --- |
|  | **Coefficient** | **Std. Err.** | **z** | **P > \|z\|** | **95% CI (L)** | **95% CI (U)** |
| **Intercept** | 0.334 | 0.350 | 0.955 | 0.339 | -0.351 | 1.019 |
| **Stage[T.AS]** | 0.053 | 0.085 | 0.629 | 0.529 | -0.112 | 0.219 |
| **Stage[T.QS]** | -0.052 | 0.085 | -0.614 | 0.539 | -0.217 | 0.114 |
| **Stage[T.QW]** | -0.062 | 0.085 | -0.736 | 0.462 | -0.228 | 0.103 |
| **Sex[T.M]** | -0.063 | 0.064 | -0.990 | 0.322 | -0.189 | 0.062 |
| **PMA** | -0.011 | 0.031 | -0.362 | 0.717 | -0.072 | 0.049 |
| **Group Var.** | 0.591 | 0.958 |  |  |  |  |
| **b) Dependent Variable: BiS** |  |  |  |  |  |  |
|  | **Coefficient** | **Std. Err.** | **z** | **P > \|z\|** | **95% CI (L)** | **95% CI (U)** |
| **Intercept\*** | -0.722 | 0.253 | -2.854 | 0.004 | -1.218 | -0.226 |
| **Stage[T.AS]** | 0.165 | 0.146 | 1.128 | 0.259 | -0.121 | 0.451 |
| **Stage[T.QS]\*** | 1.132 | 0.146 | 7.755 | < 0.001 | 0.846 | 1.418 |
| **Stage[T.QW]\*** | 0.682 | 0.146 | 4.675 | < 0.001 | 0.396 | 0.969 |
| **Sex[T.M]** | 0.162 | 0.111 | 1.467 | 0.142 | -0.055 | 0.379 |
| **PMA\*** | -0.193 | 0.053 | -3.627 | < 0.001 | -0.298 | -0.089 |
| **Group Var.** | 0.259 | 0.254 |  |  |  |  |
| **c) Dependent Variable: local nPAC** |  |  |  |  |  |  |
|  | **Coefficient** | **Std. Err.** | **z** | **P > \|z\|** | **95% CI (L)** | **95% CI (U)** |
| **Intercept\*** | -0.261 | 0.101 | -2.594 | 0.009 | -0.459 | -0.064 |
| **Stage[T.AS]** | 0.013 | 0.080 | 0.165 | 0.869 | -0.144 | 0.170 |
| **Stage[T.QS]\*** | 0.457 | 0.080 | 5.709 | < 0.001 | 0.300 | 0.613 |
| **Stage[T.QW]** | -0.097 | 0.080 | -1.207 | 0.227 | -0.253 | 0.060 |
| **Sex[T.M]\*** | 0.147 | 0.061 | 2.422 | 0.015 | 0.028 | 0.266 |
| **PMA\*** | -0.221 | 0.029 | -7.571 | < 0.001 | -0.278 | -0.164 |
| **Group Var.** | 0.032 | 0.062 |  |  |  |  |

For wPLI, no significant effects were found for PMA (β = -0.011, *p* = 0.717), behavioral state, or sex, suggesting that phase synchronization remained relatively stable across maturation and behavioral states (Tab. 1a). In contrast, BiS showed a significant negative association with PMA (β = -0.193, *p* < 0.001), indicating a decrease in bistability with cortical maturation. Similarly, local nPAC exhibited a strong negative correlation with PMA (β = -0.221, *p* < 0.001), suggesting a developmental reduction in phase-amplitude coupling. Compared to SOAS, QS showed significantly higher nPAC (β = 0.457, *p* < 0.001), while AS (β = 0.013, *p* = 0.869) and QW (β = -0.097, *p* = 0.227) did not differ significantly (Tab. 1c). Additionally, male infants exhibited significantly higher nPAC than females (β = 0.147, *p* = 0.015), while no sex differences were observed for BiS or wPLI. We observed broad-band effect of such negative correlations for both BiS and local nPAC (Fig.4). Maximal negative correlation for BiS occurs in θ band during QW and AS, while it extends to faster rhythms in SOAS, and it remains confined to δ band during QS (Fig. 4b). For local PAC, correlation with PMA was more evident between the phase of slow oscillations up to δ-band (0.2–2 Hz) and amplitude of faster oscillations up to β across all behavioral states (Fig. 4c).

**Figure 4:**
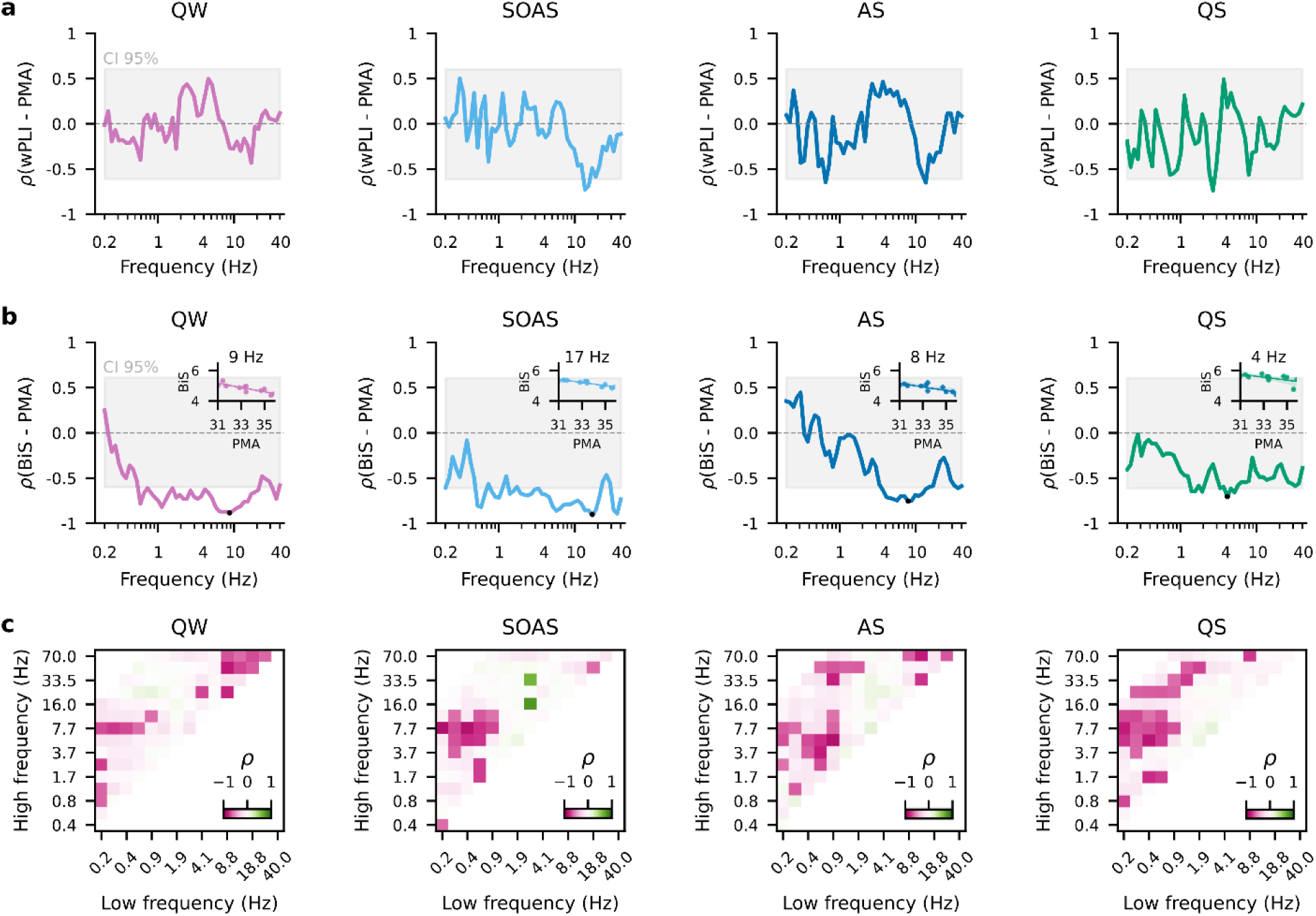
Neuronal dynamics correlate with age in specific behavioral states. **(a)** Spectra of Spearman correlation coefficient between weighted phase-lag index (wPLI) and postmenstrual age (ρ(wPLI–PMA)) and **(b)** bistability index (BiS) and postmenstrual age (ρ(BiS–PMA)), separately for each vigilance state. Shaded areas represent the 2.5^th^ and 97.5^th^ percentiles of 1’000 surrogates. Negative correlations indicate that the metric decreases with age, as illustrated in each small panel in **(b)**, where the minimum BiS-PMA correlation was chosen as an example for each behavioral state. **(c)** Spearman correlation coefficient (ρ) between local normalized phase-amplitude coupling (nPAC) and postmenstrual age for each low frequency-high frequency pair and behavioral state. Regions with less transparency correspond to areas with BH corrected *p* < 0.05.

In summary, these findings suggest that while wPLI remains stable, BiS and nPAC decrease with maturation, with notable state-dependent differences in both metrics.

## Discussion

Our study demonstrated that key electrophysiological markers (phase synchronization, bistability, and phase-amplitude coupling) follow distinct spatial and temporal trajectories during early cortical maturation in preterm infants. Bistability and phase-amplitude coupling decreased with increasing postmenstrual age (PMA), reflecting a shift from discontinuous, burst-driven dynamics toward more continuous and autonomous oscillatory patterns characteristic of maturing cortical networks. Vigilance states profoundly modulated network dynamics: quiet sleep (QS) was associated with the highest bistability and δ-band synchronization, while wakefulness featured enhanced β-to-γ coupling.

Furthermore, spatial gradients emerged, with frontal regions displaying higher bistability compared to posterior regions, aligning with known maturation patterns of cortical development.

### Vigilance states modulate cortical dynamics in preterm infants

EEG activity varied markedly across vigilance states. Quiet wakefulness (QW) exhibited elevated broadband power, particularly in the β–low γ range (Fig. 2a), and increased bistability (Fig. 2c). While these features could reflect heightened sensorimotor processing and immature state differentiation (Norman et al. 2008; Saby and Marshall 2012; Wang et al. 2023), the presence of slow oscillations (< 1Hz) during QW (Fig. 2a), combined with the rarity of this state at this age (Uccella et al. 2025), suggests that QW likely represents transitional or drowsy states under elevated sleep pressure rather than fully alert wakefulness.

By contrast, QS emerged as a distinct and highly organized brain state characterized by low power, high bistability, and strong δ-band synchronization. This pattern aligns with previous studies describing discontinuous thalamocortical activity (González et al. 2011; Tokariev et al. 2012) and large-scale slow oscillations during sleep (Vanhatalo and Kaila 2006). Therefore, QS appears to represent a state of transient network disconnection interspersed with synchronized bursts, a pattern characteristic of early cortical activity and widely implicated in shaping synaptic connectivity and supporting developmental plasticity (Steriade et al. 1993; McCormick and Bal 1997; Khazipov and Luhmann 2006; Colonnese and Khazipov 2012).

Importantly, our findings challenge the view of QS as a merely passive state. Instead, QS was defined by active oscillatory organization, with slow rhythms orchestrating the temporal structure of higher-frequency activity. PAC profiles, particularly δ-to-higher-frequency coupling, reinforced this view and point to QS as a privileged window for network reorganization during early development.

Active sleep (AS) and sleep onset active sleep (SOAS) showed reduced bistability compared to QS and pronounced θ-band synchronization (Fig. 2). This pattern resembles the spectral signature of adult REM sleep (Moroni et al. 2007; Boyce et al. 2016) and may represent early developmental precursors of REM (Roffwarg et al. 1966; de Groot et al. 2024), potentially supporting processes such as memory consolidation (Moroni et al. 2007; Boyce et al. 2016).

In addition to global and regional phase-amplitude coupling (PAC) trends, our analysis of coupling patterns revealed important state-dependent differences in the hierarchical organization of cortical coordination. During QS, PAC was predominantly driven by slow oscillations modulating the amplitude of higher-frequency activities (δ-to-β/γ), underscoring the orchestrating role of slow rhythms during early brain development activity (Dreyfus-Brisac and Monod 1965; Curzi-Dascalova 1992; Vanhatalo et al. 2005; Tolonen et al. 2007). In contrast, QW was characterized by enhanced fast-to-fast coupling (*e.g.*, β phase modulating γ amplitude), suggesting that local cortical circuits were more actively engaged and regulated in this wake-like state. Both AS and SOAS exhibited an intermediate profile, with both slow-to-fast and fast-to-fast PAC present but at lower levels. These findings highlight a dynamic reorganization of cortical coupling across vigilance states, reflecting developmental and functional shifts from global network coordination during sleep to more local, fast-frequency interactions during wakefulness.

### Regional differences in synchronization, bistability, and PAC

Beyond state-dependent modulation, clear regional differences emerged. Frontal derivations exhibited higher bistability of δ oscillations across all states, in line with the posterior-to-anterior gradient of cortical maturation (Ruoss et al. 2001; Ramenghi et al. 2007; Kurth et al. 2012; van den Heuvel and Thomason 2016). This pattern likely reflects the persistence of immature, burst-like activity in prefrontal areas. Furthermore, the frontal predominance of δ-band bistability likely mirrors the role of delta brushes and slow oscillations in orchestrating frontal network development (Arichi et al. 2017; Whitehead et al. 2017).

Phase synchronization displayed dynamic patterns across vigilance states. During QS, δ-band synchronization was globally elevated, suggesting uniform thalamocortical drive. On the other hand, during more active states (QW, AS, and SOAS), posterior regions exhibited stronger slow-frequency synchronization, possibly reflecting more active sensory processing in maturing visual and sensorimotor cortices (Blumberg et al. 2022). PAC profiles revealed regionally heterogeneous trajectories (Fig. 3c). Although statistical significance was not reached for anterior-posterior PAC differences following FDR correction, our analyses indicated stronger slow-to-fast coupling in posterior regions during QW. On the other hand, PAC from frontal regions showed higher β-to-γ PAC in QW and state-dependent slow waves to θ/σ range modulation.

### Maturational trajectories and developmental shifts in criticality

A central finding of our study is that bistability and PAC declined systematically with advancing PMA. This trajectory likely reflects a gradual reorganization of cortical network dynamics, through which initially rigid and globally synchronized network states are progressively replaced by more independent and adaptable patterns of oscillatory coordination. This interpretation is supported by recent studies spanning different stages of early brain development (Yrjölä et al. 2024). In very premature infants (24-27 weeks gestational age), emerging phase-amplitude coupling has been observed between slow waves and nested theta oscillations, reflecting an endogenous, sensory-independent mechanism thought to support the initial wiring of perisylvian networks (Moghimi et al. 2020). Later in development, global neuronal coupling modes, including amplitude-amplitude correlations and distant PAC, have been shown to decline with increasing conceptional age, indicating reduced globally coherent bursting and increasing local autonomy of cortical networks (Yrjölä et al. 2024).

Accordingly, our data indicate that, in our cohort of preterm infants born very low birth weight, local PAC remains prominent but shows a systematic decline with advancing PMA, alongside dynamic modulation by vigilance states and regional differentiation. This pattern reflects a transitional stage between globally coupled, burst-dominated dynamics and the emergence of more autonomous and flexible oscillatory regimes. However, as highlighted by the observational and correlational nature of our and previous studies (Moghimi et al. 2020), mechanistic conclusions remain inherently limited. Establishing causal relationships between these coupling modes and cortical maturation will require complementary approaches, including experimental perturbations and multimodal neuroimaging strategies.

Our findings align with and significantly extend the brain criticality hypothesis, which posits that healthy neural systems operate near a critical regime balancing order and disorder (Palva and Palva 2018; Zimmern 2020). The gradual decline in bistability and PAC that we observed with advancing PMA likely reflects a developmental tuning process that progressively guides cortical networks toward this critical regime. Early in development, the preterm brain appears to operate in a relatively supercritical state, characterized by hypersynchronization, elevated burst propensity, and strong hierarchical coupling (Hartley et al. 2012; Iyer et al. 2015). Such dynamics, while seemingly inefficient from the perspective of mature network function, may be essential during early life to drive activity-dependent wiring and synaptic refinement across widespread circuits.

As maturation progresses, the observed decline in PAC, particularly slow-to-fast coupling, and reduced bistability suggest a shift toward more distributed and autonomous oscillatory regimes. This transition may serve to prevent excessive excitation and allow the brain to achieve a dynamic balance where oscillations can flexibly couple and decouple in response to environmental demands. In this sense, our data suggest that the developing brain undergoes a form of “critical tuning”, through which early hypersynchronous states evolve into more context-sensitive and efficient modes of operation. Importantly, the state-dependent and regionally differentiated patterns observed in our cohort highlight that criticality is not a global property achieved uniformly across the cortex, but rather a multifaceted and spatially heterogeneous process.

Taken together, these findings position criticality not as a static target of brain maturation, but as a dynamic developmental trajectory involving progressive shifts in local and global network properties. Disruption or deviation from this trajectory, either through premature transition to hypoconnected states or failure to downregulate hypersynchronous dynamics, may have important implications for neurodevelopmental outcomes in preterm populations. Future studies integrating multimodal imaging, computational modeling, and longitudinal clinical follow-up will be crucial to further elucidate how critical tuning processes support or impair emerging cognitive, sensory, and motor functions.

### Conclusions and future directions

In summary, our results provide a comprehensive account of how vigilance states, cortical regions, and age interact to shape early cortical dynamics in preterm infants. Decreasing bistability and PAC reflect the gradual emergence of autonomous cortical activity, while regional differences align with known maturation gradients. Moreover, state-dependent reorganization of PAC patterns, from slow-dominated during sleep to fast-to-fast coupling during wakefulness, highlights the functional adaptation of oscillatory coordination during early life. Future research with larger samples and high-density recordings is needed to validate and expand on these findings. Ultimately, these quantitative EEG metrics may prove useful not only for describing brain maturation but also for predicting neurodevelopmental outcomes and guiding early personalized interventions.

## Abbreviations

AS: active sleep
SOAS: sleep onset active sleep
QS: quiet sleep
QW: quiet wakefulness
BiS: bistability index
EEG: electroencephalography
PAC: phase-amplitude coupling
PSG: polysomnography
PMA: postmenstrual age
PSD: power spectral density
VLBW: very low birth weight
wPLI: weighted phase-lag index

